# A single-cell transcriptomic atlas of the periventricular proliferative zone in the late gestation fetal brain in the pigtail macaque

**DOI:** 10.64898/2026.01.08.698393

**Authors:** John A. Cornelius, Megana Shivakumar, Taeyun Kim, Orlando Cervantes, Raj P. Kapur, Hazel Huang, Hong Zhao, Briana M. Del Rosario, Amanda Li, Richard J. Li, Sidney Sun, Andrew E. Vo, Gygeria Manuel, Akshaj Bharadwaj, Jeff Munson, Miranda Li, Edmunda Li, Austyn Orvis, Michelle Coleman, Alexandra Christodolou, Melissa R. Berg, Britni C. Curtis, Brenna A. Menz, Jin Dai, Inah D. Golez, Solomon N. Wangari, Stephen A. McCartney, Kristen Noble, Pilar Flores-Espinoza, Addy Cecilia Helguera Repetto, Andrea Olmos-Ortiz, Monica S. Fonseca Perez, Chris English, Audrey Baldessari, Nardhy Gomez-Lopez, Rebecca Hodge, Lakshmi Rajagopal, Kristina M. Adams Waldorf

## Abstract

**Background:** The fetal brain undergoes rapid changes in late gestation, when waves of neurogenesis and gliogenesis shape cortical circuitry. The periventricular proliferative region and adjacent white matter are enriched in neuroprogenitor cells, newborn neurons, and interneurons, which is challenging to study in the late gestation human fetal brain. The nonhuman primate (NHP) provides a powerful translational model to overcome this limitation, given its close similarity to human neurodevelopmental trajectories. The study objective was to construct a single-cell RNA-Seq (scRNA-Seq) atlas of the late-gestation fetal brain of the pigtail macaque (Macaca nemestrina), focused on the periventricular proliferative zone.

**Methods:** A sample of the lateral ventricular wall, subventricular zone, and overlying white/gray matter was dissociated into single cells and processed through the 10X Genomics pipeline, followed by SoupX removal of ambient RNA, and Seurat’s pipeline to aggregate, cluster and annotate single-cell populations. Monocle3 was used to determine pseudotime and map lineage progression.

**Results:** This analysis captured diverse populations of neuroprogenitors, newborn neurons, developing lineages of excitatory and inhibitory neurons, oligodendrocyte and astrocyte lineages, and resident immune and endothelial cells.

**Conclusions:** Single-cell populations from the third-trimester nonhuman primate fetal brain are highly similar to those in the human fetus. This late-gestation single-cell atlas of the periventricular proliferative zone provides a unique reference for progenitor, neuronal, glial, vascular, and immune cell states during a critical window of primate neurodevelopment, enabling mechanistic interrogation of how inflammatory, infectious, or hypoxic insults disrupt vulnerable neurogenic niches.

## Background

The fetal brain is a dynamic organ that undergoes rapid cellular proliferation, differentiation, and migration to establish complex cortical circuitry and complex neurologic functions. In the developing fetal brain, the ventricular zone (VZ), subventricular zone (SVZ), periventricular white matter (PVWM), and deep white matter (DWM) represent critical regions enriched in neuroprogenitor cells and newborn neurons. These neurogenic niches are highly vulnerable to injury from pathogens and inflammatory insults that access the cerebrospinal fluid (CSF), such as Zika virus (ZIKV), cytomegalovirus (CMV), and diverse bacterial infections. In addition to congenital infections, the periventricular proliferative zone is highly sensitive to sterile inflammatory exposures, hypoxia–ischemia, and preterm birth–associated immune activation, all of which converge on these neurogenic niches during late gestation. Experimental models, such as nonhuman primates (NHPs), are required to study these events because normal human fetal brain tissue at this late gestational stage is rarely accessible for ethical reasons.

NHPs provide a powerful model for human neurodevelopment due to similarities in gestational length, cortical structure, and timing of neurogenesis. Among the most used species are rhesus macaques (*Macaca mulatta*), pigtail macaques (*Macaca nemestrina*), and cynomolgus macaques (*Macaca fascicularis*), which have been utilized to study viral neuropathogenesis^1–4^, neuroprogenitor and white matter injury^3,5–7^, and neurodevelopmental outcomes after in utero exposures^8^. Although NHP models are widely employed to study how experimental insults affect the developing brain, the extent to which they recapitulate the diversity of human neural progenitors, glial precursors, and newborn neurons has been incompletely defined. While there are differences in the species-specific rates of neurogenesis and gliogenesis, the structure and organization of the late-gestation macaque brain closely resemble those of the human brain; there is a similar laminar organization of the SVZ and subplate, and parallel timelines of oligodendrocyte maturation and axonal growth.^9–13^ NHPs are one of the most translationally relevant animal models for investigating human fetal neurodevelopment and its susceptibility to infection and injury.

Single-cell RNA sequencing (scRNA-Seq) has transformed the study of complex tissues by enabling detailed investigation of cellular heterogeneity and transcriptional programs. Several scRNA-Seq atlases of the human brain have been generated, primarily from mid-gestation samples (gestational weeks 15–24), providing critical insights into progenitor populations, lineage trajectories, and neuronal subtype specification.^14^ However, human datasets from late gestation are extremely limited due to limited tissue availability. In contrast, scRNA-Seq studies of the NHP fetal brain are rare, with only a handful of reports describing bulk or spatial transcriptomics, organoids, or specific progenitor cell populations.^2,15,16^

Our objective was to construct a scRNA-Seq atlas of the late gestation pigtail macaque fetal brain, focusing on cell populations in the periventricular proliferative zone and the adjacent white and gray matter, which are vulnerable to infection or inflammatory injury due to exposure to cerebrospinal fluid. By characterizing progenitor, neuronal, glial, and immune cell populations at single-cell resolution, we provide a reference dataset for the normal development of these vulnerable regions. This atlas also establishes a foundation for comparative studies with human fetal brain datasets and provides a resource for investigating how congenital infections or inflammatory exposures alter neurodevelopmental trajectories.

## Results

### Study Design

To determine the single-cell transcriptional profiles of the fetal pigtail macaque within the periventricular proliferative zone and adjacent white/gray matter, we collected tissues from these regions for single-cell dissociation and analysis using the 10X Genomics pipeline. Tissues were obtained from 9 healthy, uninfected fetal pigtail macaques, serving as historical controls for infectious disease studies. They had either undergone surgical catheterization and saline inoculation of the amniotic cavity and choriodecidual space (N=4) or subcutaneous media inoculation of the forearm (N=5; **Table 1**). All animals were delivered by Cesarean section, and none developed preterm labor or other perinatal complications. Fetal brain tissues from the periventricular zone and adjacent white/gray matter were single-cell dissociated and processed through the 10X Genomics scRNA-Seq pipeline and integrated using the Seurat analysis pipeline.

**Table 1.**
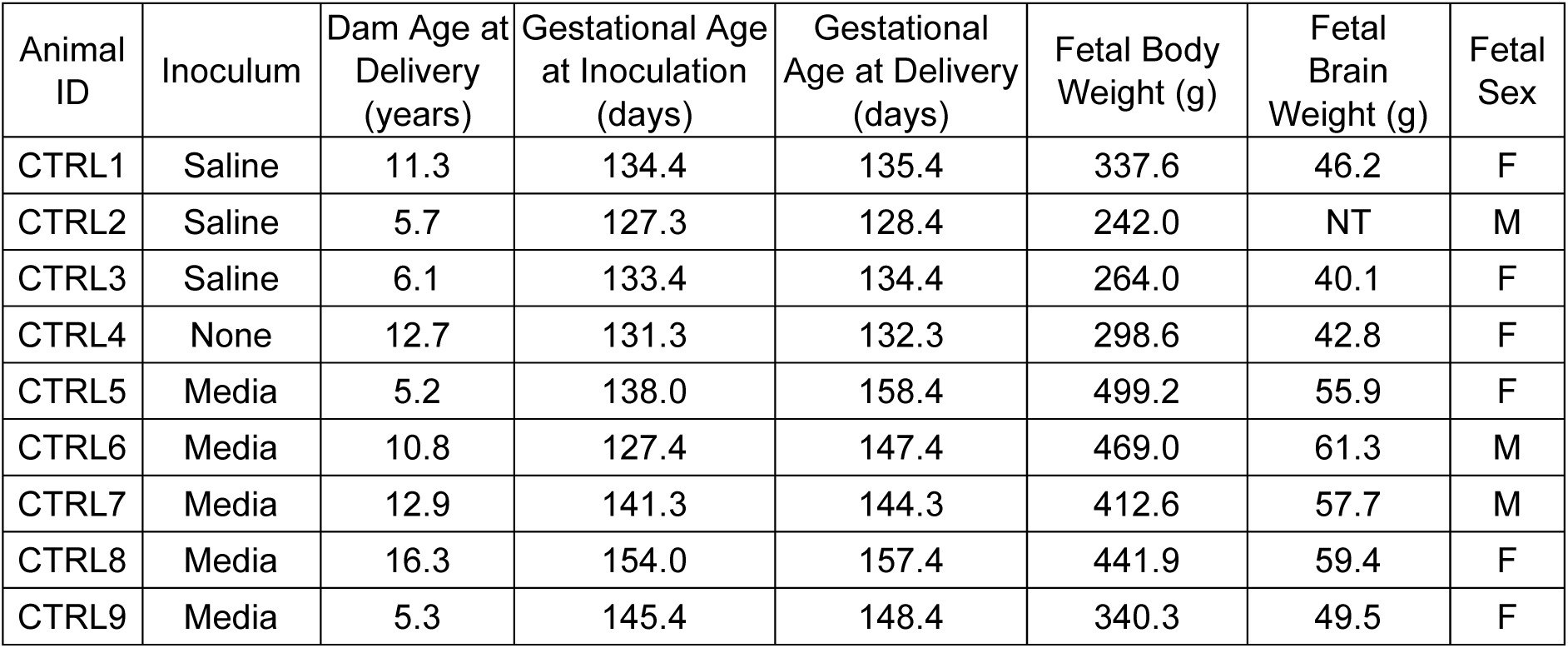
This table shows the inoculum, maternal age at delivery, gestational age at inoculation and delivery, fetal body weight, fetal brain weight, and fetal sex for each animal in the study. Abbreviations: F, female; g, grams; M, male; and NT, not tested/weighed.

### Comparative Anatomy of the Late Gestation Fetal Pigtail Macaque and Early Infant Human Fetal SVZ and Deep White Matter

Creation of a scRNA-Seq atlas of the fetal pigtail macaque SVZ and deep white matter necessitates an understanding of anatomical similarities and differences with human fetuses at the same time point (**Fig. 1**). As normal third-trimester human fetal brains are rarely obtained and described, we compared the same anatomical structures with an early infant brain. Overall, the architecture of the fetal SVZ and deep white matter is remarkably similar between the two species.

**Figure 1.**
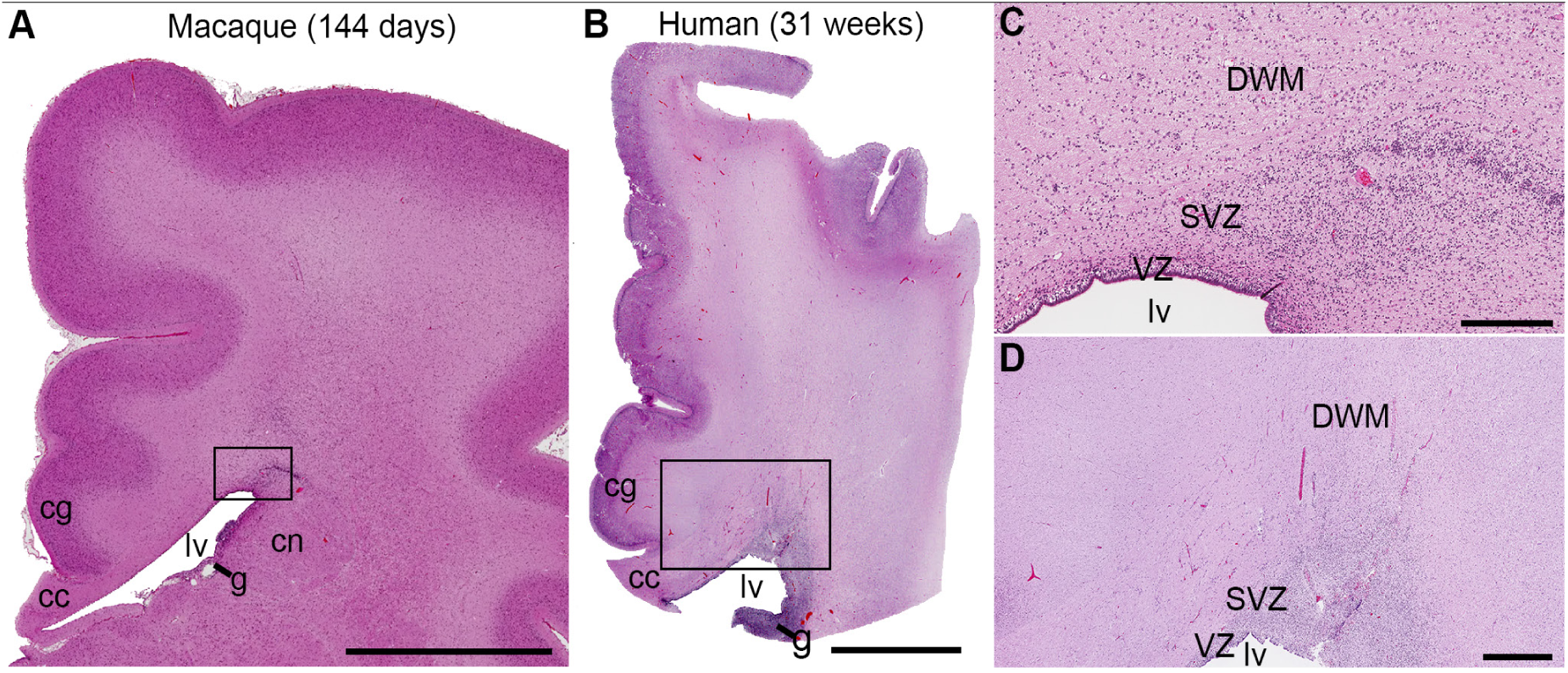
Histology of cerebral cortical histology in a pigtail macaque and human third-trimester fetuses at comparable ages. This figure shows images of the cerebral cortex from pigtail macaque and human fetuses at comparable gestational ages in the third trimester (macaque: 144 days, term gestation: 172 days; human: 31 weeks, term gestation 40 weeks). (A) A low magnification image of a hematoxylin-and-eosin-stained coronal section through the dorsal portion of a macaque cerebral hemisphere includes the lateral ventricle (lv) with adjacent ganglionic eminence (g), corpus callosum (cc) and caudate nucleus (cn). (B) The medial half of a similar section from a human comparable stage human fetus shows many of the same landmarks, although the human brain is significantly larger. (C) Higher magnification of the rectangular region outlined in (A) illustrates relative positions of the ventricular zone (VZ), subventricular zone (SVZ), and deep white matter (DWM). (D) Higher magnification of the rectangular region outlined in (B) shows the locations of the same zones in the human fetus. Scale bars: A, 5 mm; B, 500 mm; C, 300 μm; D, 1 mm.

### Clustering, Annotation, and Cell Origin

Unsupervised clustering identified 27 transcriptionally distinct populations, which were manually annotated using established lineage-specific marker genes, comparative primate and human fetal brain datasets, and trajectory relationships (**Fig. 2**) Cell populations were visualized using a uniform manifold approximation and projection (UMAP) plot, and then trajectory analysis was performed to calculate pseudotime using Monocle. An undifferentiated progenitor/intermediate (UP/I) population was at the center of the UMAP with most populations radiating outward from (or inward to) the UP/I. Key populations near the UP/I radial glia (vRG), outer radial glia undergoing gliogenic differentiation (oRG-G), immature neurons (ImmN), immature astrocytes (AS0), pericytes (PC), and a population of committed oligodendrocyte progenitors (COP2). Although QC analysis was performed initially in the scRNA-Seq pipeline, we reanalyzed the final clusters to determine whether these populations contained few features or unique genes, or high mitochondrial reads. The number of genes detected per cell was high in most cell populations with a few exceptions: ImmN, oligodendrocyte 2 (OL2), and UP/I (**Fig. 3A**). Total counts of unique molecular identifiers (UMI) were also low in these populations reflecting a lower complexity and quality of these clusters (**Fig. 3B**). Average mitochondrial read percentage was below 1% across all clusters, including ORG-G (0.73%), the population with the highest mitochondrial reads (**Fig. 3C**); relatively higher mitochondrial reads are typical for a rapidly dividing glial population. Most clusters in the atlas were connected to the UP/I population, including vRG, ORG-G, ImmN, AS0, COP2, and PC. The distribution of proportions and counts for each population varied across individual animals and with gestational age (**Fig. 4).** One of the most striking temporal changes in the atlas was the appearance of COP2, OL1, and OL2 at approximately 144 days of gestation, which accompanied a decrease in the relative percentage of the OPC population. With advancing gestational age, there was also a decreasing proportion of border-associated microglia (BAM) and MG0 and an increasing proportion of MG1.

**Figure 2.**
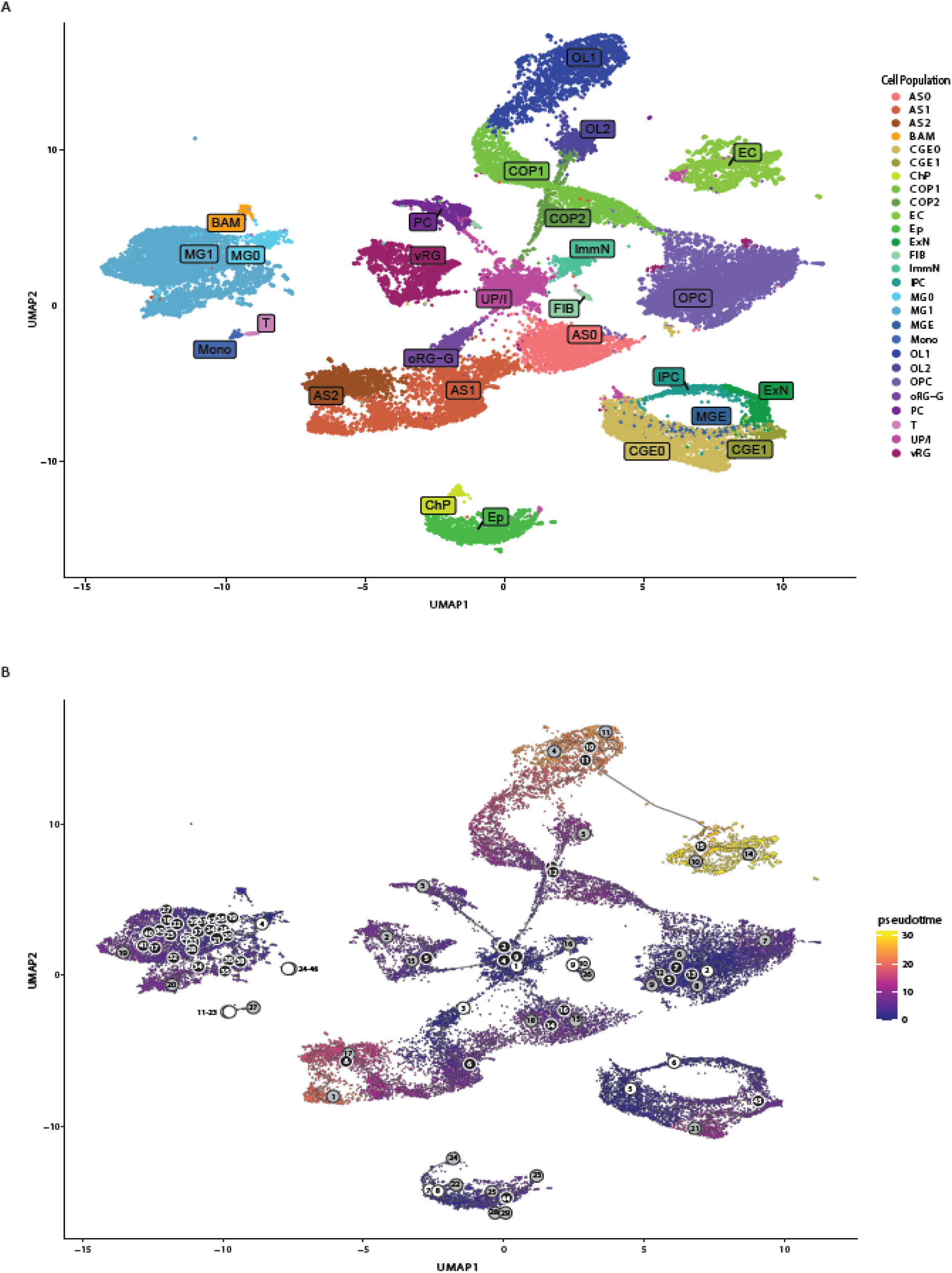
Single-Cell Atlas of the Periventricular Proliferative Region of the Late-Gestation Pigtail Macaque Fetal Brain. (A) UMAP of single-cell transcriptomes illustrate distinct cellular clusters identified by unsupervised clustering. Each point represents an individual cell, colored according to cluster (B) The trajectory plot displays inferred lineage relationships among clusters. Roots were inferred based on gene expression patterns within clusters and labeled with white-numbered circles, indicating the start of pseudotime. Branch points, labeled with black-numbered circles, represent major lineage bifurcations, while gray-numbered circles mark terminally differentiated populations along the trajectory. Abbreviations: AS, astrocyte; BAM, border-associated macrophage; T, T cells; CGE, caudal ganglionic eminence lineage; ChP, choroid plexus; COP, committed oligodendrocyte progenitor; EB, erythroblast; EC, endothelial cell; EP, ependymal cell; ExN, excitatory neuron lineage; ImmN, immature neurons; oRG-G, outer radial glia undergoing gliogenic differentation; MG, microglia; MGE, medial ganglionic eminence lineage; Mono, monocyte; OL, oligodendrocyte; OPC, oligodendrocyte progenitor cells; PC, pericyte; and vRG, ventricular radial glia

**Figure 3.**
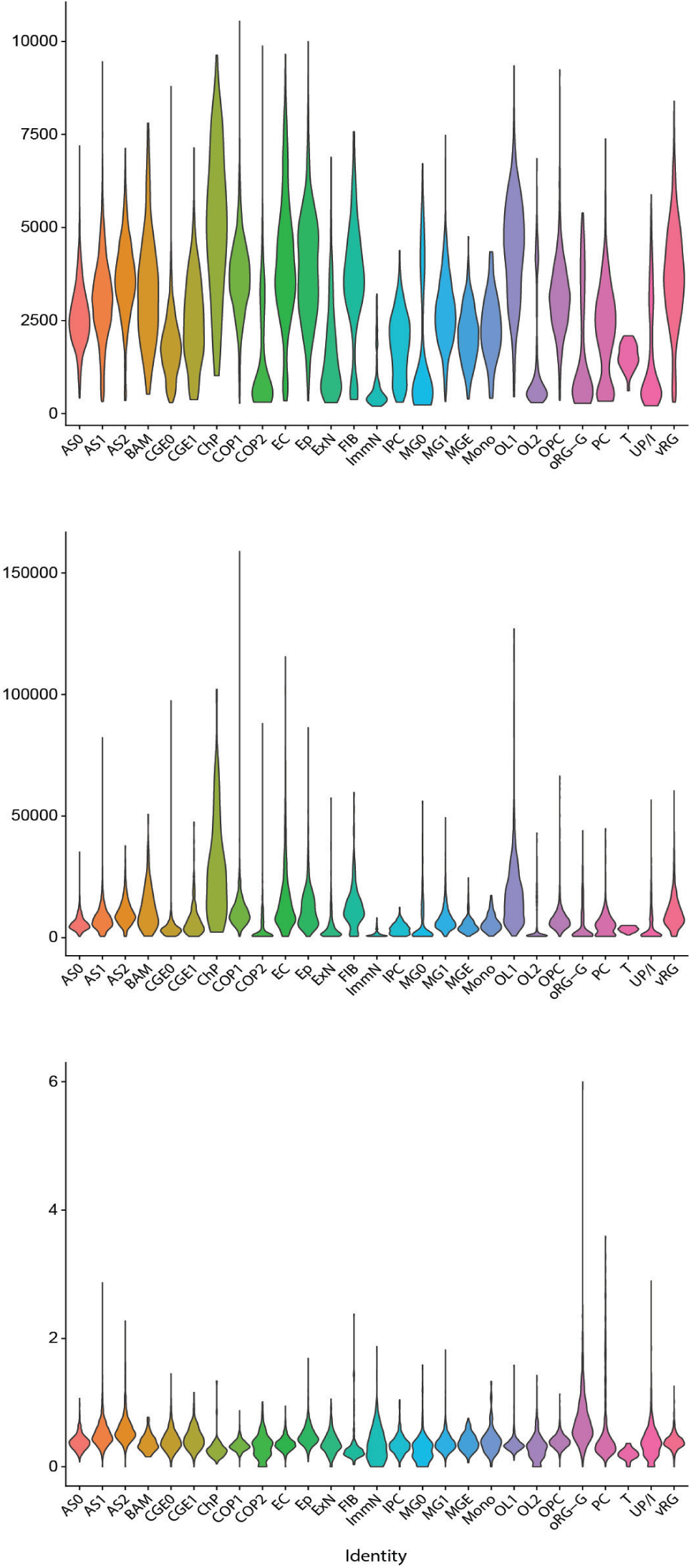
Quality Control Analyses of Cell Populations. Panels show the number of genes detected (A), total counts of unique molecular identifiers (UMI, B), and the percentage of mitochondrial reads (C). Abbreviations are in the legend of Figure 2.

**Figure 4.**
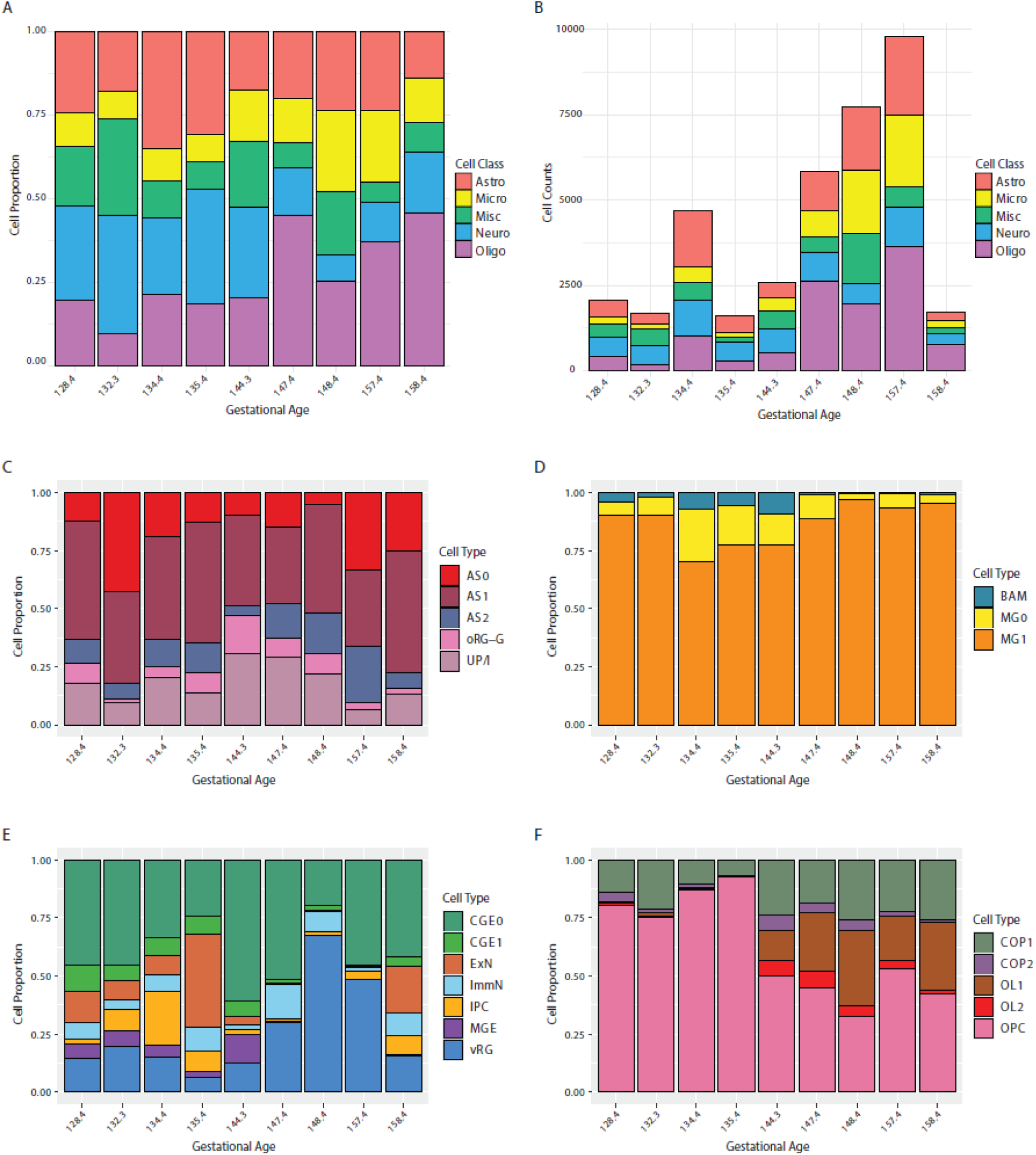
Cell Proportions and Cell Counts for Single-Cell Populations. These stacked bar plots show either the cell proportion (A, C, D, E, F) or cell counts (B) for the annotated clusters for each animal in order along the x-axis by the gestational age at necropsy. Abbreviations are found in the legend of Figure 2. Subpopulations within a single cell type were aggregated to show the changes in the proportion of cell populations (A) or counts (B) as a function of gestational age. (C-F) Subpopulations within a single cell type were grouped with similar cell types to show how cell proportion changed over gestational age for cells within the astrocyte (C), microglia (D), neuron (E), and oligodendrocyte (F) lineage. Abbreviations are in the legend of Figure 2.

### Neuronal Lineages

Three progenitor populations were annotated in our atlas: ventricular radial glia (vRG), outer radial glia (oRG-G) differentiating into a gliogenic lineage, and intermediate progenitor cells (IPC; **Fig. 5A**). vRG was identified by high expression of cell cycle genes, including MKI67, CDK1, TOP2A, PCNA, ASPM, and ANLN (**Fig. 5B**). oRG-G expressed HOPX, FAM107A, and TNC with a trajectory into the astrocyte lineage (**Fig. 2, 5B**).

**Figure 5.**
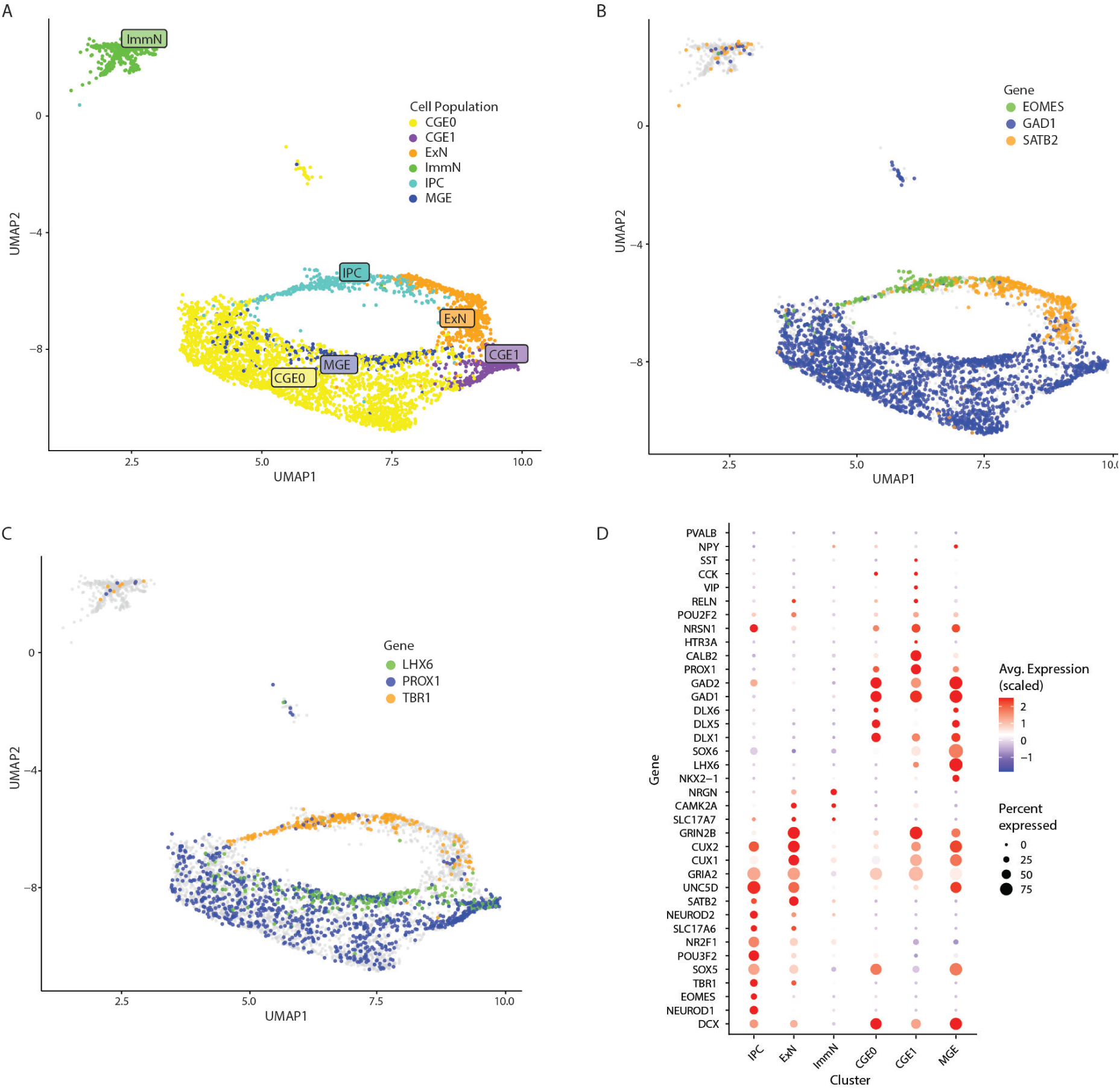
Transcriptomic characterization of neuron subpopulations. (A) UMAP visualization of neuron populations identified by single-cell RNA-Seq. Each point represents an individual cell, and colors denote cluster identity. (B) Feature plot showing expression of key transcriptional markers along the excitatory lineage: *EOMES* (green), *GAD1* (blue), and *SATB2* (orange). These markers highlight the transition from early progenitor-like to mature excitatory neuronal states. (C) A UMAP feature plot displays differential expression of LHX6, PROX1, and TBR1, defining genes for MGE, CGE, and excitatory neuron lineages. (D) A dot plot summarizes the expression of representative excitatory neuronal genes across clusters. Dot size reflects the proportion of cells expressing each gene, while color intensity indicates scaled average expression. Cell population abbreviations are defined in the Figure 2 legend.

Intermediate progenitor cells (IPCs) arising from radial glial in the periventricular proliferative zone undergo several rounds of cell division before differentiating into glutamatergic projection neurons that populate both the deep layers (DL, layers V-VI) and upper layers (UL, layers II-IV) of the neocortex. The IPC population in the periventricular proliferative zone expressed EOMES (TBR2) and TBR1, consistent with a late transitional IPC state that exited the cell cycle and is transcriptionally committed to a deep-layer neuronal fate. UMAP feature plots illustrated a progressive maturation from EOMES⁺ early neurons (IPCs) toward SATB2⁺/TBR1⁺ differentiated populations (ExN). RNA expression within the ExN population was consistent with a transitional subplate neuronal state that had initiated upper-layer and callosal transcriptional programs (CUX2, SATB2, POU3F2, NR2F2) while retaining residual deep-layer markers (TBR1, TLE4, SLC17A6). ExN also expressed SLC17A6 (VGLUT2), a canonical marker of glutamatergic neurons that is highly expressed in early born deep-layer excitatory neurons and subcortical projection neurons. In contrast, the ImmN population expressed SLC17A7 (VGLUT1), NRGN, and CAMK2A, but few other markers of mature excitatory or inhibitory neurons, suggesting an immature state.

Medial and caudal ganglionic eminences (MGE and CGE, respectively) produce GABAergic interneurons in the developing fetal NHP and human brain. These eminences form proliferative zones within the ventral telencephalon, located along the ventral and medial walls of the lateral ventricles. The CGE extends caudally and produces VIP^+^, CALB2^+^, and NR2F2^+^ interneurons that migrate into superficial cortical layers. Together, the MGE and CGE transient germinal structures generate most GABAergic inhibitory interneurons in the brain. MGE-derived interneurons were defined by NKX2-1, LHX6, DLX1, DLX2, GAD1 and GAD2 expression (**Fig. 5D**). Although the MGE gives rise to PVALB^+^ and SST^+^ interneurons and basal ganglia projection neurons, PVALB was not expressed in these clusters, indicating that these MGE interneurons were relatively immature. A population of CGE-derived progenitors, labeled CGE0, was characterized by the new-onset expression of NR2F1 (COUP-TF1), NR2F2 (COUP-TFII), DLX1, DLX5, and GAD2. Differentiation of the CGE lineage progressed through CGE1, which expressed the migration marker, NRP2. CGE2 was a more differentiated CGE interneuron population indicative of a VIP⁺/CALB2⁺ interneuron fate that expressed PROX1, CALB2, KCNC1, CHRNA2, and VIP. Expression of GABAergic markers in both the MGE and CGE lineage (GAD1, GAD2, DLX2, ERBB4) reflected their inhibitory identity (**Fig. 5D**). In summary, these data delineate transcriptionally distinct progenitor and interneuron precursor populations in the developing ventral forebrain adjacent to the subventricular zone.

### Oligodendrocyte Lineage

Five cell populations within the oligodendrocyte lineage were identified by expression of canonical markers, including SOX10, PDGFRA, PCDH17, OLIG2, BCAS1, MAL, PLP1, and MBP (**Fig. 6**). Of the oligodendrocyte lineage markers, OPC expressed only PDGFRA, indicating a population entering the oligodendrocyte program. Committed oligodendrocyte progenitors (COPs) were defined by expression of BCAS1 and ENSMMUG00000056728 (ortholog of NKX2-2), indicating a transition from OPC to an early pre-myelinating or differentiating oligodendrocyte. OL1 and OL2 expressed genes consistent with mature myelinating oligodendrocytes, including MBP, MOG, PLP1, and MAL. Together, these analyses delineate the transcriptional progression from PDGFRA⁺ oligodendrocyte progenitors through ENPP6⁺ committed precursors to MBP⁺/PLP1⁺ mature oligodendrocytes.

**Figure 6.**
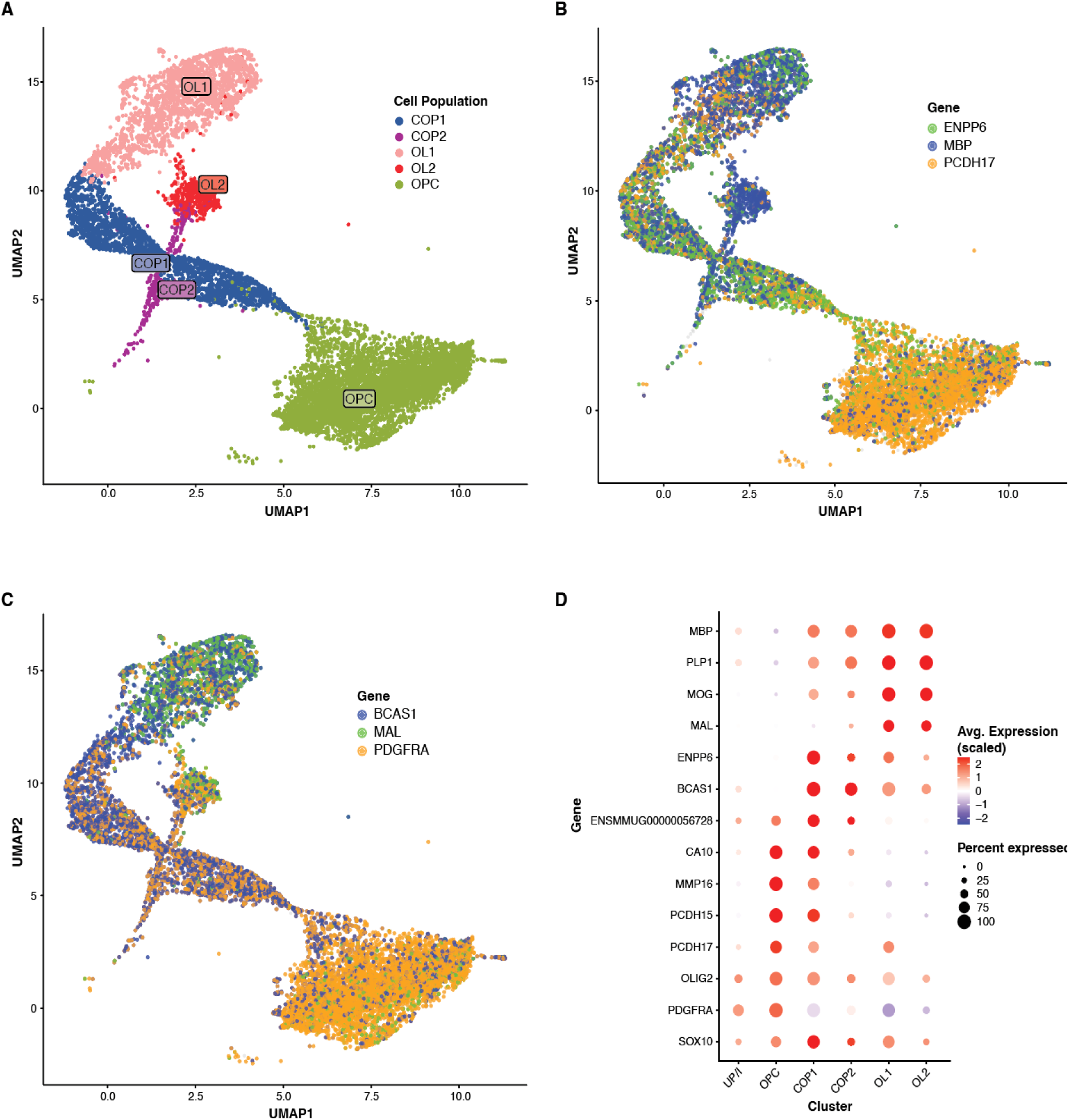
Transcriptomic landscape of oligodendrocyte lineage cells during differentiation. (A) UMAP visualization of oligodendrocyte lineage clusters identified by single-cell RNA-Seq, including oligodendrocyte progenitor cells (OPC), committed oligodendrocyte precursors (COP1–COP2), and maturing oligodendrocytes (OL1–OL2). Each point represents a single cell, and colors denote cluster identity, illustrating the continuum from progenitor to mature oligodendrocyte states. (B) Feature plots show expression of representative lineage-associated genes. (C) Expression of additional oligodendrocyte lineage markers. (D) A dot plot summarizing gene expression of key oligodendrocyte genes across clusters. Dot size indicates the percentage of cells expressing each gene, and color intensity represents scaled average expression levels. Cell population abbreviations are in Figure 2.

### Astrocyte Lineage

Astroglial populations clustered into three populations (**Fig. 7A**) along a trajectory of pseudotime (**Fig. 7B**). The earliest population was labeled as AS0 and had low expression of SLC1A3 (EEAT1), a canonical astrocyte marker as well as low expression of SLC1A2 the first glutamate transporter induced during gliogenesis, and NFIA, a master regulator of astrocyte gene expression programs (**Fig. 7B**). The AS1 population was marked by higher expression of these markers as well as high expression of TNC, HOPX, and TIMP3. AS2 was the most mature astrocyte population and expressed GFAP and FAM107A, consistent with acquisition of homeostatic functions. An alternative differentiation pathway was observed starting in the oRG-G population feeding into AS1. The co-expression of several canonical astrocyte markers is shown in **Fig. 7C-F**. Together, these data reveal transcriptionally distinct astrocyte subtypes that vary in expression of key structural, metabolic, and signaling genes, reflecting molecular diversity within the developing astroglial lineage.

**Figure 7.**
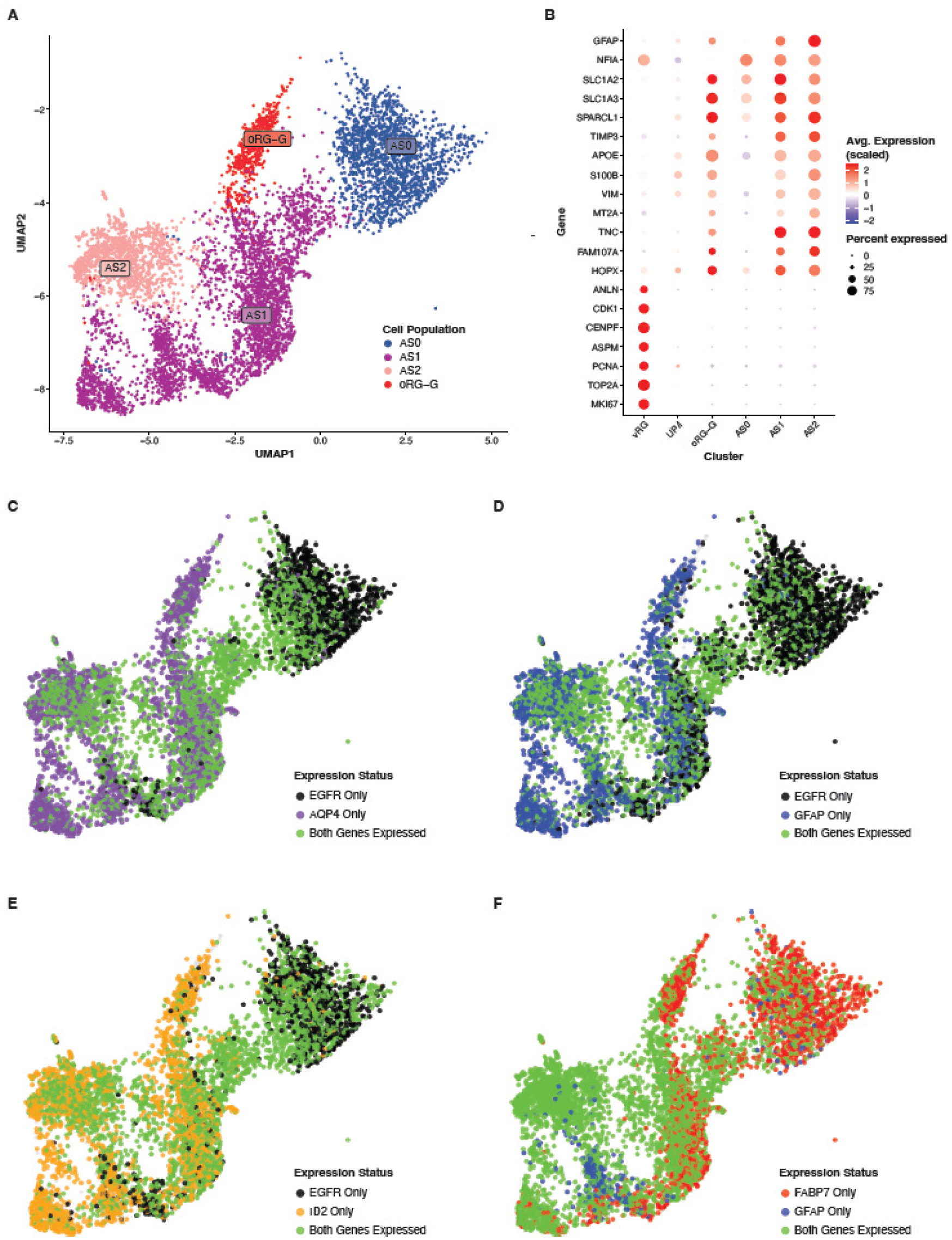
Transcriptional profiling of astrocyte subpopulations. (A) UMAP visualization of astrocyte clusters (AS0–AS2) and oRG-G identified by single-cell RNA-Seq. Each point represents an individual cell, and colors denote distinct astrocyte subpopulations, illustrating transcriptional heterogeneity among astroglial cells. (B) A dot plot is shown to summarize expression of representative astrocyte-enriched genes. Dot size indicates the percentage of cells expressing each gene, and color intensity represents scaled average expression levels. (C-F) Co-expression feature plots are shown. Cell population abbreviations are in Figure 2.

### Microglia

Microglial populations exhibited transcriptional features of late fetal maturation, characterized by expression of AIF1 (IBA1), CX3CR1, P2RY12, CSF1R, and TREM2, denoting a homeostatic and surveillant phenotype. Unsupervised hierarchical modeling generated 2 distinct clusters, with IL1B and CD86 being key differentiators between these two populations (**Fig. 8A**). AIF1 expression in these clusters was constitutive (**Fig. 8B**). BAM was characterized by high expression of CD14, MRC1, CD86, and CD4; this combination of markers is most characteristic of macrophages that tend to reside in the choroid plexus or ventricular surfaces, locations that were enriched in this data (Fig. 8C)

**Figure 8.**
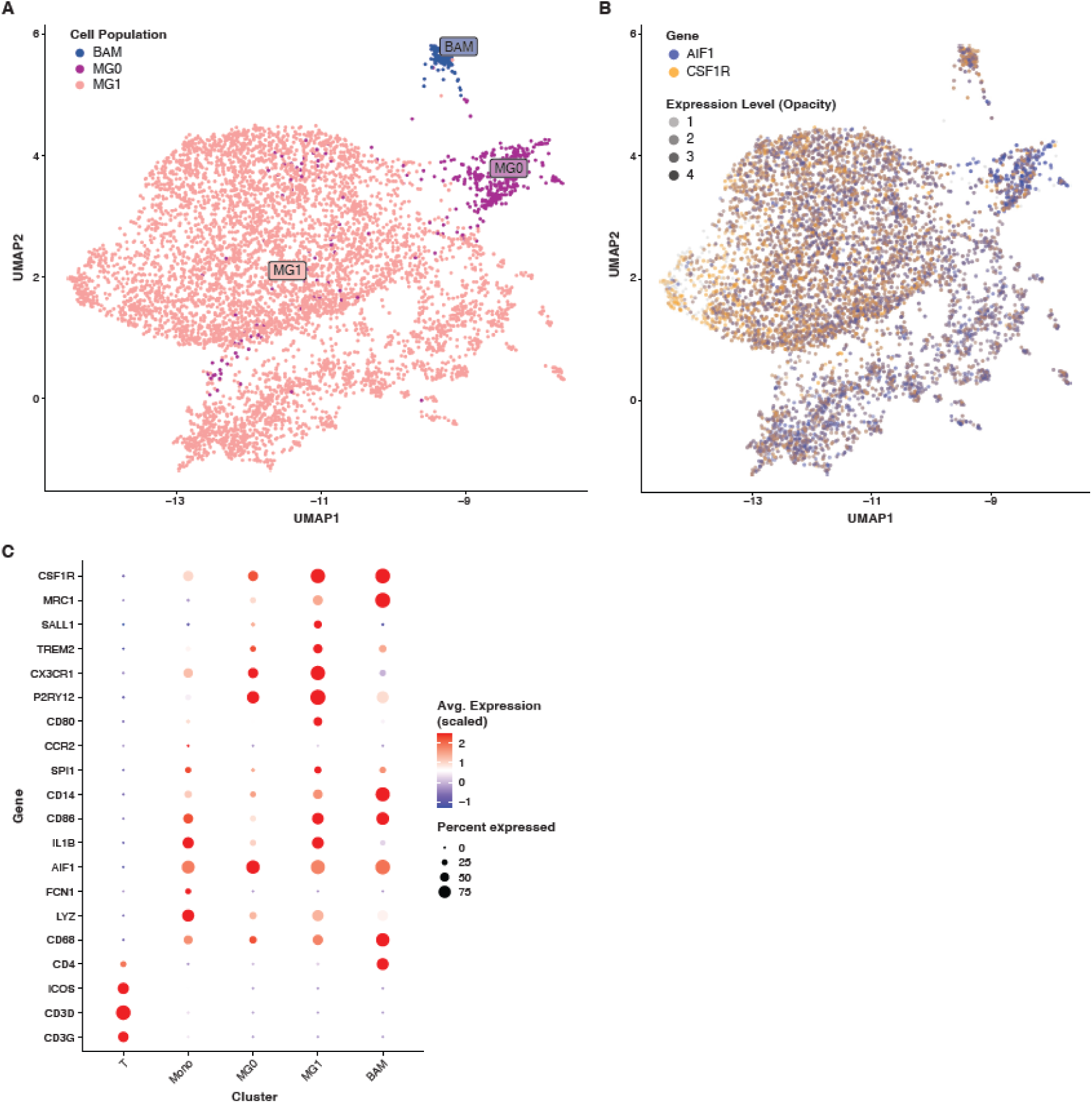
Transcriptional profiling of microglial subpopulations. (A) UMAP visualization of microglial clusters (MG0, MG1 and BAM) identified by single-cell RNA-Seq. Each point represents an individual cell, and colors denote distinct microglial subpopulations, illustrating transcriptional heterogeneity among microglial cells. (B) A feature plot is shown for canonical microglial markers AIF1 and CSF1R, confirming microglial identity and regional or maturation-associated differences in gene expression across clusters. (C) A dot plot is shown to summarize expression of representative microglia-enriched genes. Dot size indicates the percentage of cells expressing each gene, and color intensity represents scaled average expression levels. Cell population abbreviations are in Figure 2.

### Stromal, Vascular, and Epithelial Cell Populations

Several other non-neuronal cell populations were identified, which would be expected in the developing fetal brain (**Fig. 9**). Pericytes (PC) ensheathe capillaries and help to stabilize capillary networks. We identified a single pericyte population that expressed PDGFRB, MYOF, ABCC9, and TAGLN. The endothelial cell (EC) population expressed endothelial cell canonical markers: FLT1, CD34, PECAM1, and VWF. Expression of AQP4, CETN2, DNALI1, and RFX2/3 characterized ependymal cells. T cells (T) were identified by CD3D, CD3G, ICOS, and CD4 expression (**Fig. 7C**). Monocytes (Mono) were identified expressing high levels of LYZ, IL1B, and CD86. The choroid plexus (ChP) was identified by its expression of TTR, CLDN3/4, and FOXJ1. COL1A2, VIM, DCN, and FMOD expression identified a small population of Fibroblasts (FIB).

**Figure 9.**
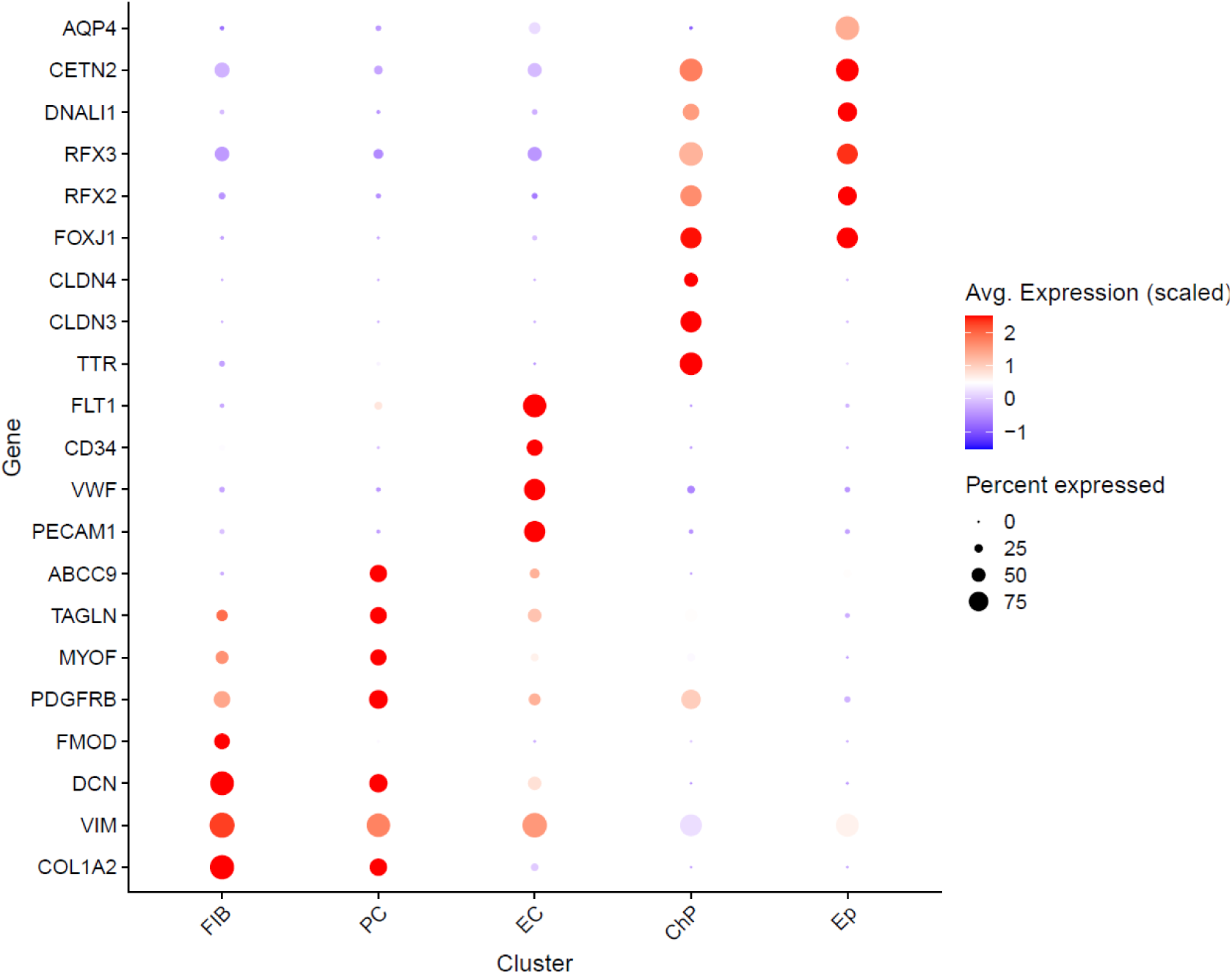
Canonical Gene Markers for Stromal, Vascular, and Epithelial Cell Populations. Canonical gene markers for fibroblasts (FIB), pericytes (PC), endothelial cells (EC), choroid plexus (CP), and ependymal cell (Ep) populations.

### Expression of Viral Entry Receptors

As many viruses are known to infect the fetal brain, we investigated the RNA expression of well-established or putative viral entry receptors, co-factors, or enzymes associated with congenital and neonatal neuroinvasive infectious diseases (**Fig. 10-11**). We focused on genes that were highly upregulated in specific fetal brain cell types, reasoning that these expression patterns might reveal cellular vulnerabilities to viral infection. Notably, a small population of border-associated microglia in the late-gestation fetal brain expressed at least one viral entry receptor exploited by multiple neurotropic viruses, including Japanese encephalitis virus (JEV: TFRC/CD71), Zika virus (ZIKV: HAVCR1/TIM-1, AXL), varicella-zoster virus (M6PR), human cytomegalovirus (CMV: NRP2), human immunodeficiency virus (HIV: CD4), and Nipah virus (NiV: EFNB2). Several other viral entry genes were upregulated in neuroprogenitors or immature neurons: LRP8 in IPCs (tick-borne encephalitis virus, TEV), CXADR in CGE1, MGE, and IPCs (coxsackievirus B viruses, CVB), and PDGFRA in vRG and UP/I (CMV). Immune cells were targets of several viruses with high IGF2R expression in monocytes (VZV) and CD4/CXCR4 co-expression in T cells (HIV). In the case of human CMV, multiple cell types in the late-gestation fetal brain may be permissive to infection. Although predicting viral infection solely from RNA expression is challenging, this resource provides insight into which cell types may be key targets.

**Figure 10.**
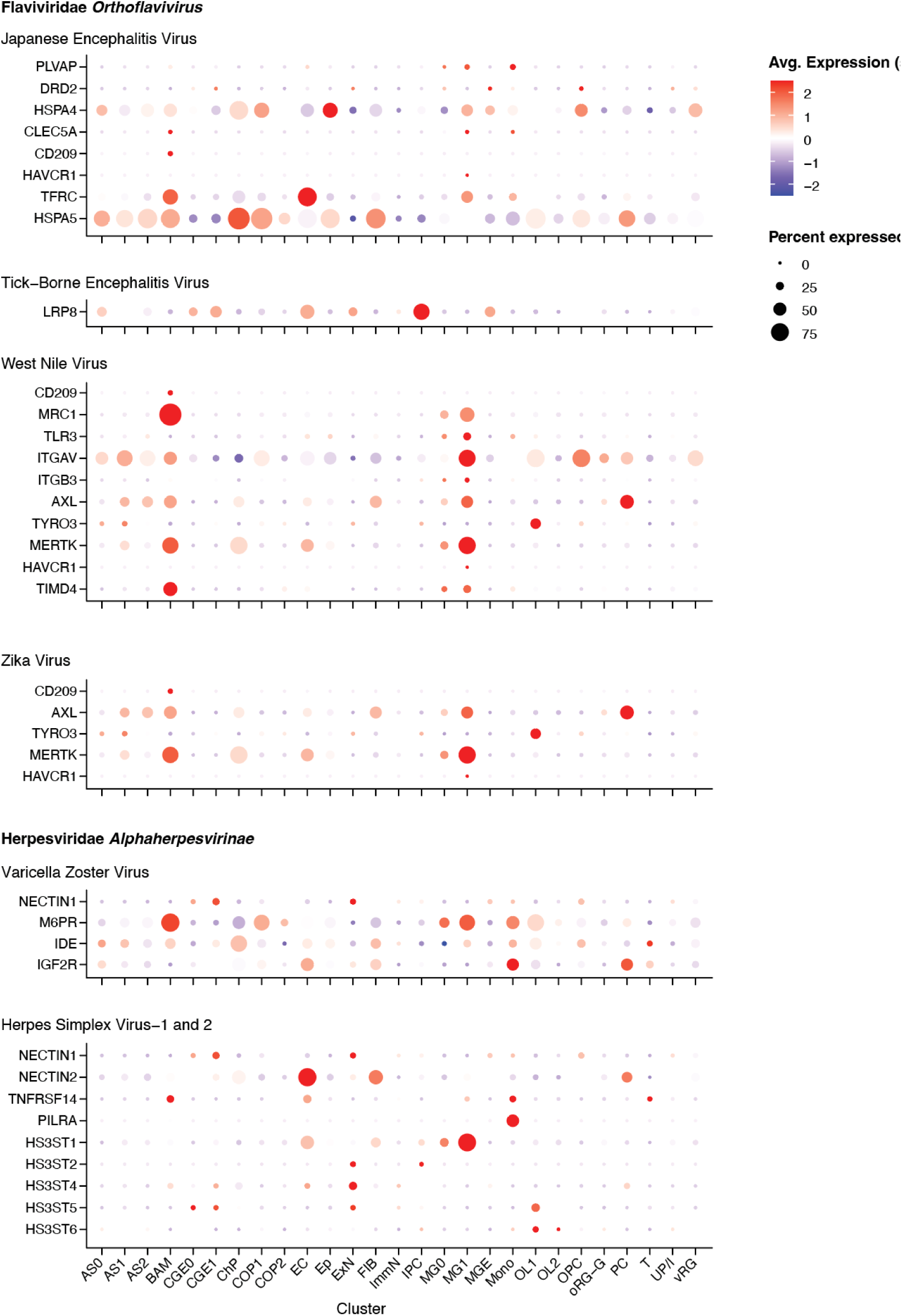
Gene Expression of Viral Entry Receptors, Co-Factors, and Enzymes in the Flaviviridae *Orthoflavivirus* and Herpesviridae *Alphaherpesvirinae* Families in the Fetal Periventricular Proliferative Region. Genes for established or putative viral entry receptors, co-factors, or enzymes are shown for select viruses associated with neuroinvasive congenital or neonatal disease in these families.

**Figure 11.**
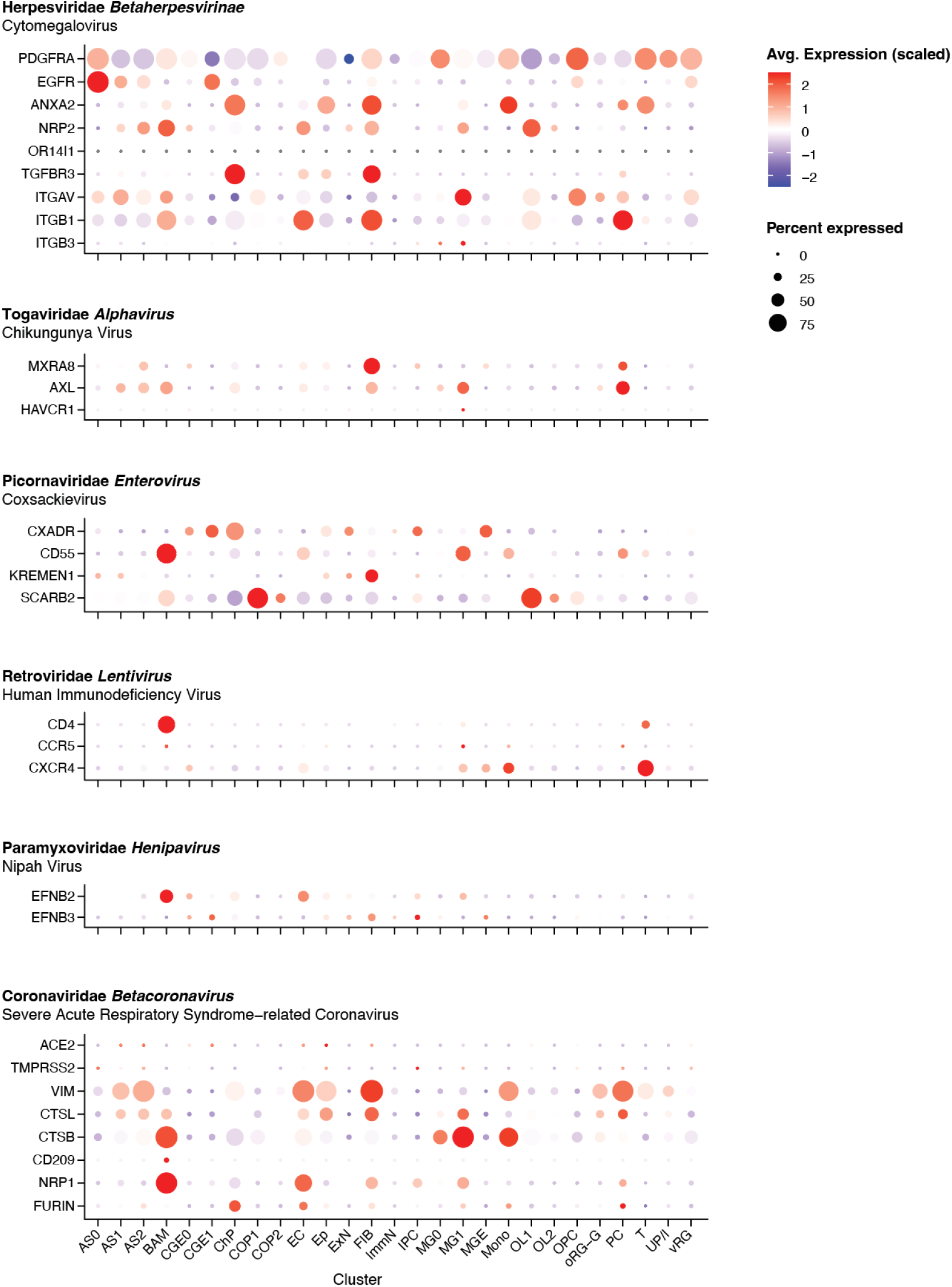
Gene Expression of Viral Entry Receptors, Co-Factors, and Enzymes of Diverse Viruses in the Fetal Periventricular Proliferative Region. Genes for established or putative viral entry receptors, co-factors, or enzymes are shown for select viruses associated with neuroinvasive congenital or neonatal disease in these families.

## Discussion

This scRNA-Seq atlas of the periventricular proliferative zone in the late-gestation NHP brain serves as a critical reference for defining normative progenitor, neuronal, and glial populations, in a region of the fetal brain that is vulnerable to injury from preterm birth, hypoxia-ischemia, mechanical ventilation, inflammatory stimuli, and congenital infections.^2,17–26^ The atlas revealed a high degree of similarity in the cellular populations of the periventricular proliferative region with those of humans and identified maturing neuronal and glial lineages, alongside smaller populations of endothelial, fibroblast, choroid plexus, ependymal, and immune cells. A scRNA-Seq reference atlas to describe normal transcriptional programs in the periventricular region can also provide insight into how hypoxic, inflammatory, and other insults disrupt neuroprogenitor development and lead to ventriculomegaly, periventricular leukomalacia, and/or long-term neurodevelopmental deficits.^27,28^

Similarities in the single-cell populations between the macaque and human brain in late gestation are expected; both human and nonhuman primate brains are characterized by early symmetric divisions of neural progenitor that expand the progenitor pool, followed by asymmetric divisions that generate neurons and glia organized into cortical columns.^29–34^ Radial glia scaffold the migration of cortical neurons during a prolonged prenatal period of neurogenesis and neuronal maturation, resulting in the complex, gyrencephalic brain characteristic of primates^33^, which contrasts with the lissencephalic brains of other species. In the late-gestation fetal macaque brain, we identified key subpopulations of radial glia and neural progenitors, including actively cycling vRG (MKI67+); oRG-g (HOPX+), associated with the astrocyte lineage; IPC (EOMES+), in a trajectory toward ExN differentiation; and the UP/I population in an intermediate progenitor state positioned downstream of vRG and upstream of oRG-g and IPC. At this time in gestation, neurons remained relatively immature with poor expression of markers characteristic of mature interneurons (MGE: PVALB, SST; CGE: VIP, RELN).

Cluster analysis revealed heterogeneity in glial populations. Within the astrocyte lineage, the ORG-g occupied the earliest pseudotime and expressed HOPX, consistent with an immature or progenitor-like transcriptional state. AS0 was characterized by high EGFR, but low GFAP expression, consistent with a developmentally immature state, and was closely associated with the UP/I. AS1 was positioned between the oRG-g and AS0 populations on the UMAP, suggesting an intermediate state between two potential astrocyte precursors. In the oligodendrocyte lineage, we identified a canonical pathway in which OPCs transitioned through a committed oligodendrocyte precursor (COP1) and matured into oligodendrocytes (OL1). A second trajectory originated from the UP/I population, bypassing the OPC stage, converging at the COP stage, and producing a distinct oligodendrocyte population (OL2). Whether OL1 and OL2 represent subpopulations in distinct spatial niches (e.g., ventricular/subventricular versus intermediate zone) or were produced in different temporal waves is unknown. Consistent with prior developmental studies, astrocyte and oligodendrocyte lineages in our dataset display partially redundant transcriptional programs, with subtle distinctions driven by cellular origin, niche, or timing of differentation.^35–45^

There are several advantages of this late gestation scRNA-Seq atlas of the periventricular proliferative region over other human and NHP atlases. Although human fetal brain atlases provide broader anatomical coverage, they have been mainly restricted to early gestation^46–49^, used a spatial^46^ or single-nucleus RNA-Seq (snRNA-Seq) approach^50^, or focused on other areas of the brain^51^. When third-trimester human fetal brains have been studied, the tissue is collected post-mortem and analyzed using spatial or snRNA-Seq. Two rhesus macaque fetal brain atlas have been described, but these were collected from the first half of gestation and focused on progenitor and neuron lineages.^52,53^ Other atlases have used combinations of fetal brain explants and human induced pluripotent stem cell-derived mixed neural cultures to define pathways induced during ex vivo stimulation with Zika virus.^54^ Mouse scRNA-Seq atlases of the fetal brain also exist^55^, but murine neurodevelopment occurs over a shorter timeline than in humans and nonhuman primates. Therefore, the nonhuman primate is a powerful model of human brain development because it shares fundamental aspects of cortical lamination, progenitor biology, and an extended gestational maturation that closely mirror human neurodevelopment.

This single-cell RNA sequencing atlas provides an important resource for identifying fetal brain cell types that may be vulnerable to neuroinvasive viruses associated with congenital and neonatal infections. For viruses that exploit multiple entry pathways, such as human cytomegalovirus, our data suggest that numerous cell types in the late-gestation fetal brain may be permissive to infection. Notably, border-associated microglia appear particularly susceptible to a broad range of viruses; their strategic localization at fetal brain barrier regions suggests they may serve as an early cellular interface facilitating productive infection and neuroinvasion. In our late-gestation atlas, a key viral entry receptor for ZIKV (AXL) was upregulated in microglia, BAM, and astrocytes, but not vRG, a key target population in the fetal brain; dynamic changes in vRG expression over gestation may explain reduced AXL expression in vRGs, and potentially, a lower susceptibility to ZIKV infection in the third trimester.^56,57^ More broadly, this atlas enables systematic screening for viral entry receptors in the fetal brain associated with emerging viruses that could cause congenital infection. Finally, a notable strength of this atlas is the inclusion of immune and border-associated macrophage populations, which are often underrepresented in human fetal brain datasets but play critical roles in shaping neurodevelopmental responses to inflammatory and infectious exposures.

The use of the pigtail macaque, whose gestational timeline and placentation parallel human development^58–60^, further enhances the translational relevance of this atlas for modeling neurodevelopmental disease and congenital infection. While histopathologic and imaging approaches have described the gross architecture of periventricular injury, they cannot resolve the cellular subpopulations affected or the mechanisms by which transcriptional programs are perturbed. This scRNA-Seq atlas provides a resource to overcome this limitation by identifying normal cellular transcriptional states in late gestation and by uncovering transcriptional networks that shape local immune and microglial activation. A limitation of this study is the collection of tissue from only the late second and third trimesters, which provides a temporal snapshot from late in gestation but leaves earlier neurogenic waves unexplored. This scRNA-Seq atlas is also focused on the periventricular proliferative zone and does not profile other important brain regions. There are also several species differences between macaque and human neurodevelopment to consider: slower postsynaptic density protein accumulation in the human fetal brain and distinct postnatal trajectories for specific lineages and regions after birth.^61, 62^ Finally, while scRNA-Seq defines cellular diversity at high resolution, functional validation through *in situ* hybridization, spatial transcriptomics, and lineage tracing is important to confirm inferred trajectories.

In summary, this late-gestation scRNA-Seq atlas of the periventricular proliferative region provides a transcriptional map of a highly vulnerable area of the developing primate brain. The late second and third trimesters represent a critical window of human brain development, during which the periventricular proliferative zone and adjacent intermediate and white matter regions undergo rapid expansion, gliogenesis, and maturation of cortical circuitry. Maternal infectious exposures and immune activation have been linked to fetal brain injury using murine and NHP models, as well as through epidemiologic studies of children.^63–68^ Fetal exposure to a viral mimetic [poly(I:C), polyriboinosinic–polyribocytidylic acid) during pregnancy has been associated with disruptions in neuroprogenitor proliferation and migration, altered cortical layering and glial organization, brain growth, and deficits in attention and behavior in offspring.^69–76^ This atlas enables comparison of normal and perturbed developmental trajectories in NHP models, and these insights may point to pathways that could be leveraged for neuroprotection or repair. An interesting finding in our atlas is that glial maturation is supported by redundant transcriptional programs, in which both canonical and alternative transcriptomic pathways independently reach mature oligodendrocyte and astrocyte states. Future studies using this atlas as a benchmark can determine how prenatal insults of varying types (e.g., inflammation, infection, preterm birth) perturb single-cell populations in this critical area of the fetal brain.

## Methods

### Ethics Statement

All nonhuman primate experiments were performed in strict accordance with the recommendations in the Guide for the Care and Use of Laboratory Animals of the National Research Council and the Weatherall report, “The use of non-human primates in research.” The Institutional Animal Care and Use Committee (IACUC) of the University of Washington (UW) approved the study’s protocols (#4165-01, last approval date: 05/07/2025; #4165-02, last approval date: 04/09/2025). All surgeries were performed under general anesthesia, and all efforts were made to minimize pain and distress.

### Experimental design

Nine healthy, pregnant pigtail macaques were enrolled as uninfected controls for two different experimental protocols. One group received choriodecidual and intra-amniotic saline infusions after surgical implantation of choriodecidual and amniotic catheters, which were compared to experimental infections with Group B Streptococcus (N=4). A second group received five subcutaneous inoculations of media along the forearm to serve as controls for a study of Zika virus pathogenesis (N=5). Gestational ages at tissue collection spanned the late second and third trimester (118-150 days of gestation; term gestation = 172 days). After delivery by cesarean section, tissues from the ventricular zone, subventricular zone, periventricular white matter, and deep white matter were collected from the fetal brain. Samples were collected in Hibernate-E medium (Thermo-Fisher, A1247601) on wet ice, and single-cell dissociation was performed within 1 hour after collection.

### Single Cell RNA-Seq and Bioinformatics Pipeline

#### Single Cell Dissociation

The right hemisphere of each brain was sectioned coronally at approximately 0.5 cm intervals. From the coronal sections, brain samples for scRNA-Seq were manually removed to prioritize sampling of the lateral ventricular wall (ventricular zone, VZ), subventricular zone (SVZ), and adjacent white matter. This protocol evolved over the course of the study. Initially, only the periventricular region of the parietal cortex was selectively sampled, but we obtained insufficient cells to complete the scRNA-Seq protocol. Therefore, we shifted to sampling full-thickness portions of the parietal cortex, including the lateral ventricle wall, VZ, SVZ, deep white matter, and adjacent white/cortical gray matter. Later, we carved out the lateral ventricle wall, VZ, SVZ, and deep white matter from the frontal, parietal, temporal, and occipital coronal sections, excluding cortical gray matter and deep nuclei. Most samples were obtained using the latter approach.

#### Isolation of Cells from the Fetal Brain

Fetal brain tissues were placed into cold HBSS with calcium and magnesium (Thermo-Fisher, 14025092) and finely chopped into approximately 0.2 mm³ cubes using sterile scissors. Tissue fragments were enzymatically digested in 2 mL of a papain digestion buffer. The digestion buffer consisted of 2.5 U/mL papain (Worthington #LS003126), 250 U/mL DNase I (Qiagen, #79254), and 10 mM HEPES (Thermo Fisher, #15630130), prepared in HBSS without calcium or magnesium (Thermo Fisher, #14175095). Digestion was carried out at 37 °C for 20 minutes, and the tube was gently inverted every 5 minutes to mix.

Following digestion, the digestion buffer was quenched with 6 mL of cold quenching buffer. L-15 media consisting of 440 mL Leibovitz L-15 media (Thermo-Fisher, #21083027), 53 mL purified H2O, 5 mL of 1M HEPES (Thermo Fisher, **#**15630130), and 0.4% BSA (Fisher Scientific, **#**50-121-5315) was mixed with 250 U/mL DNase I (Qiagen, #79254) and 0.5 mg/mL trypsin inhibitor (Sigma-Aldrich, T2011) to make the quenching buffer. Following quenching and centrifugation at 200 x g for 5 minutes at room temperature, the supernatant was removed, and an additional 1 mL of quenching buffer was added. Using a P-1000 micropipette, the sample was gently pipetted up and down to disperse the tissue into a single cell suspension after approximately 15 cycles.

To wash the sample, the cells were transferred to a new 50 mL tube and resuspended in 40 mL of resuspension buffer, prepared with the same materials as the quenching buffer, but without trypsin inhibitor. The suspension was then filtered with a 40-μm cell strainer (Fisher Scientific, #22-363-547) and centrifuged at 200 x g for 10 minutes at room temperature. The supernatant was removed, and the cells were resuspended to 10 mL of resuspension buffer. Cells were then counted, and cell viability was assessed using trypan blue (Fisher Scientific, #MT25900CI). Single-cell libraries were prepared immediately following dissociation.

#### Sequencing

10x Chromium single-cell libraries were prepared according to the standard protocol outlined in the manual (Chromium Next GEM Single Cell 3’ kits v3.1: dual index, 10X Genomics, Pleasanton, CA). Briefly, we loaded single-cell suspensions, 10x barcoded gel beads, and reagents onto a “Chromium Single Cell A Chip” to generate Gel Bead-In-EMulsions (GEMs) using the 10X Chromium Controller (10X Genomics, Pleasanton, CA). Reverse transcription of mRNA occurred inside each GEM, followed by amplification of cDNA and preparation of sequencing libraries according to manufacturer instructions. Quality control was performed in multiple steps to check the quality of the cDNA and the library using the Agilent 2200 TapeStation system (Agilent Technologies, Santa Clara, CA). Finally, shallow sequencing was performed using a NextSeq 2000 (Illumina Inc., San Diego, CA) to evaluate the percentage of reads mapping to the reference genome, the distribution of GC content, and sequencing depth (number of reads per cell). Based on this information, we determined if the cDNA library was of sufficient quality to send for deep sequencing and whether an adjustment to the amount of any samples was necessary to achieve a greater number of reads per cell. Deep sequencing of the cDNA library was performed on a NovaSeq (Illumina Inc., San Diego, CA) to achieve approximately 40,000 reads per cell.

### Genome Alignment

Raw FASTQ files were processed using Cell Ranger v7.2.0 software (10x Genomics, Pleasanton, CA, USA). Reads were aligned to the rhesus macaque genome (Mmul_10, GenBank Assembly ID: GCA_003339765.3), which has a contiguous assembly and is more completely annotated than the pigtail macaque genome. Because the rhesus and pigtail macaque are closely related species with high conservation of coding and regulatory regions, alignment to the rhesus macaque reference genome enabled robust transcript detection while minimizing gene annotation loss that would occur using the less complete pigtail macaque assembly.

### Quality Control, Normalization, and Harmonization

To reduce ambient RNA contamination in our data, we applied the SoupX package.^77^ Raw (unfiltered) and filtered gene-cell count matrices generated by Cell Ranger were imported into SoupX, which estimates the profile of ambient RNA (“soup”) from empty droplets and models its contribution to each captured cell. The contamination fraction was estimated using the autoEstCont function with default parameters and adjusted where necessary based on expression of known cell-type-specific marker genes. Corrected count matrices were generated using adjustCounts and subsequently used for downstream quality control, normalization, clustering, and differential gene-expression analyses. Empty droplets and potential doublets were removed by filtering out cells with fewer than 200 genes or more than 2500 genes. Damaged cells were removed by filtering out cells with greater than 3% mitochondrial reads.

With the Seurat package in R (Seurat version 5.1.0, R version 4.4.0)^78–82^, samples were aggregated and the data set was normalized with the SCTransform^83^ function. The first 50 principal components were found using the RunPCA function with 30 selected for evaluation. Next, scDEED was used to detect dubious cell embeddings within the default UMAP settings and then to optimize UMAP parameters to minimize distortion.^84^ Data was corrected for batch effects and integrated using the Harmony package (version 1.2.0)^85^ and the top 30 harmony components were processed using the Seurat runUMAP function to embed and visualize the cells in a two-dimensional map via the Uniform Manifold Approximation and Projection for Dimension Reduction (UMAP) algorithm. A resolution of 0.25 was initially used to cluster single cells. Clusters were evaluated for distinct marker genes, and the top expressing genes ranked by log fold change (logFC) were found for each cluster using the FindTopMarkers function in Seurat. Based on this analysis, it was determined that some clusters needed to be subdivided while others were merged due to similarity in gene expression. After cluster annotation, scCustomize was then used to generate final UMAPs.^86^

### Trajectory Inference

Trajectory analyses used Monocle 3 (R) (tested on R 4.4.0; monocle3 ≥1.3.7). All code used default parameters unless otherwise stated; random seeds were fixed for reproducibility. Size-factor and dispersion normalization were performed within Monocle 3. Highly variable genes (HVGs) were imported from Seurat. Pseudotime was then computed for all cells as geodesic distance from the root along the principal graph and scaled to [0,1] for visualization. Graph branch points were identified directly from the principal graph. Lineages were assigned by following edges from the root to terminal nodes (tips) and by evaluating marker enrichment along each path.

## DECLARATIONS

### Ethics Approval

The Institutional Animal Care and Use Committee (IACUC) of the University of Washington (UW) approved the study’s protocols (#4165-01, last approval date: 05/07/2025; #4165-02, last approval date: 04/09/2025). All surgeries were performed under general anesthesia, and all efforts were made to minimize pain and distress.

### Consent for Publication

N/A

### Availability of Data and Materials

The data generated and analyzed during the current study is available in the <u>NCBI Gene Expression Omnibus (GSE305531)</u>. Bioinformatics code is available on the [Adams Waldorf Lab GitHub](https://github.com/kadamswaldorflab).

### Authors’ Contributions

KMAW supervised the research and analysis performed in this project, secured funding, analyzed the data, supervised the research team, and wrote and edited the manuscript. JC and LR also supervised aspects of this project. JC, MS, TK, AB, JM, were involved in data curation, formal analysis, and software. AL, RL, and EL were also involved in data curation. JC, MS, TK, JM, and RPK performed the data analysis and created visual representations of the data. JC, MS, TK, OC, RPK, HH, HZ, BDR, AL, RL, SS, AEV, GM, AB, JM, ML, EL, AO, MC, MB, BCC, BAM, JD, IDG, SNW, SAC, KN, PF-E, ACHR, AO-O, MSFP, CE, and AB were investigators on this project and collected data. KAW and LR secured the funding. LR also assisted in project administration. KMAW and JC wrote the first draft of the manuscript. NGL and RH assisted in data analysis and interpretation. All authors edited and revised subsequent drafts and approved the final manuscript.

## Acknowledgment

We are grateful to Riley Raker for assistance with graphic design. We thank Devesh Bajaj, Nehmat Bhutani, and Maria Waldorf for assistance with cluster annotation.

