## Supplemental Information for "A single-cell transcriptomic atlas of the periventricular proliferative zone in the late gestation fetal brain in the pigtail macaque"

### **Table S1.** Animal Demographics

| Animal ID | Inoculum | Maternal Age at Delivery (years) | Gestational Age at Inoculation (days) | Gestational Age at Delivery (days) | Fetal Body Weight (g) | Fetal Brain Weight (g) | Fetal Sex |
| --- | --- | --- | --- | --- | --- | --- | --- |
| CTRL1 | Saline | 11.3 | 134.4 | 135.4 | 337.6 | 46.2 | F |
| CTRL2 | Saline | 5.7 | 127.3 | 128.4 | 242.0 | UNK | M |
| CTRL3 | Saline | 6.1 | 133.4 | 134.4 | 264.0 | 40.1 | F |
| CTRL4 | None | 12.7 | 131.3 | 132.3 | 298.6 | 42.8 | F |
| CTRL5 | Media | 5.2 | 138.0 | 158.4 | 499.2 | 55.9 | F |
| CTRL6 | Media | 10.8 | 127.4 | 147.4 | 469.0 | 61.3 | M |
| CTRL7 | Media | 12.9 | 141.3 | 144.3 | 412.6 | 57.7 | M |
| CTRL8 | Media | 16.3 | 154.0 | 157.4 | 441.9 | 59.4 | F |
| CTRL9 | Media | 5.3 | 145.4 | 148.4 | 340.3 | 49.5 | F |

This table shows the inoculum, maternal age at delivery, gestational age at inoculation and delivery, fetal body weight, fetal brain weight, and fetal sex for each animal in the study. Abbreviations: F, female; g, grams; M, male; UNK, unknown.
